# A novel behavioral paradigm for characterizing anticipatory postural adjustments in mice

**DOI:** 10.64898/2026.11.10.765800

**Authors:** Egzona Morina, Federico Claudi, Andrew J Murray

## Abstract

Voluntary movements perturb the body’s equilibrium, and the nervous system limits this disturbance by generating anticipatory postural adjustments (APAs) that redistribute body weight before and during movement. APAs have been characterized extensively in humans and larger animal models, but the absence of a tractable behavioral assay in mice has limited access to the genetic and circuit-level tools needed to investigate their neural basis. Here, we describe a force-plate behavioral paradigm in which freely moving mice stand on four independent load cells, receive an auditory GO cue and initiate locomotion voluntarily without task-specific training. Using force recordings, high-speed video and markerless pose estimation, we show that mice initiate a step with a stereotyped sequence of weight shifts: the center of pressure (CoP) moves anteriorly and then laterally away from the swing side, the swing forepaw and diagonally opposite hindpaw unload, and body weight is carried by the remaining diagonal pair. A single principal component explained approximately 80% of the variance in the average trial, compared with approximately 50% when paw identity was shuffled. Pressure recordings identified rapid swing-forepaw unloading beginning, on average, 87.9 ms before lift-off. Head and ear movement began approximately 200 ms before lift-off, and an operational multimodal estimate placed the interval between the earliest detected postural movement and the onset of forepaw rotation at approximately 160-180 ms. Neuropixels recordings from the pontine reticular nucleus (PRN) and supplementary motor area (SMA), monosynaptic tracing of inputs to reticulospinal PRN neurons, and preliminary cell-ablation experiments identify the PRN as a candidate node in the circuitry supporting anticipatory postural control. This paradigm provides a platform for circuit-level investigation of postural control and its disruption in disease.

## Introduction

Everyday voluntary movements such as stepping, reaching or rising from a chair displace the body’s center of mass relative to its base of support. They are nonetheless executed fluidly and without falling, because the nervous system generates postural adjustments in advance of, and during, the movement itself. These anticipatory postural adjustments (APAs) are commonly understood as the motor read-out of a feedforward process: the brain predicts the mechanical consequences of the upcoming movement and pre-emptively scales muscle activity in the trunk and limbs to inimize the resulting disturbance (Belenkii et al., 1967; Bouisset & Zattara, 1981; Massion, 1992; Aruin & Latash, 1995). In human gait initiation, anticipatory postural adjustments help establish the forward velocity reached by the end of the first step (Lepers & Brenière, 1995).

APAs are typically divided into two phases (Schepens & Drew, 2003). Preparatory APAs (pAPAs) begin before any visible movement of the limb and create the initial imbalance, and momentum, required to initiate the action. They end when the limb first leaves the ground. Accompanying APAs (aAPAs) continue throughout the movement and keep the body in dynamic equilibrium until it is completed. In humans, gait initiation is preceded by anticipatory changes in muscle activity and center of pressure that help generate forward movement and prepare the transfer of support between the legs (Lepers & Brenière, 1995; Mizusawa et al., 2017). Postural control also adapts to prior experience and environmental context. This is illustrated by the “broken escalator” phenomenon, in which individuals show a forward-sway aftereffect when stepping onto a stationary platform previously experienced as moving, despite knowing that it is stationary (Bronstein et al., 2009). Anticipatory postural control is affected by aging, with older adults showing delayed anticipatory muscle responses to predictable perturbations (Kanekar & Aruin, 2014). People with Parkinson’s disease also exhibit reduced APA amplitudes and altered timing during step initiation (Jacobs et al., 2009).

Studies in quadrupeds have used the same logic. Cats and dogs trained to lift a limb or reach into a tube show a stereotyped diagonal weight-redistribution pattern before the limb leaves the ground, in which the CoP shifts away from the limb to be lifted and body weight is carried on the remaining paws (Coulmance et al., 1979; Gahery et al., 1980; Ioffe et al., 1982; Schepens & Drew, 2003). In cats, neurons in the pontomedullary reticular formation, the origin of the reticulospinal tract, are active during these postural adjustments (Prentice & Drew, 2001; Schepens & Drew, 2004; Schepens et al., 2008). Reticulospinal neurons are well placed to contribute: they receive convergent input from many brain regions and project rapidly to spinal circuits controlling axial and limb muscles (Sivertsen et al., 2014; Brownstone & Chopek, 2018). Cortical areas, notably the supplementary motor area (SMA), also shape APA timing, as shown in patients with Parkinson’s disease (Jacobs et al., 2009).

Despite this, the circuit mechanisms that generate APAs remain poorly understood. Human experiments cannot readily manipulate defined cell types, and work in cats and primates is constrained by the available tools. Mice offer cell-type-specific genetic access, viral tracing, large-scale electrophysiology and disease models, but no behavioral paradigm for measuring APAs in mice has been available. Here we report such a paradigm. It measures the weight borne by each paw during unconstrained step initiation, requires no task training, and is compatible with high-speed video and neural recording. We use it to describe mouse APAs and compare them with those of other species, and we combine it with neural recordings, circuit tracing and cell ablation to ask whether the pontine reticular nucleus (PRN) is a plausible node in the circuit that generates them.

## Results

### A force-plate paradigm for freely moving mice

Measuring APAs in a quadruped requires (i) a platform that permits natural locomotion, (ii) sensors able to resolve the weight borne by each of the four paws before and during a step, and (iii) an event that defines the start of the motor plan so that pAPAs and aAPAs can be separated using limb movement. We met these requirements with four single-point load cells arranged beneath the paws of a standing mouse (Fig. 1A; separation of the front paws 1.8 cm, of the hind paws 2.0 cm, front-to-hind 3.5 cm, scaled to the average mouse body length of ∼8 cm).

**Figure 1.**
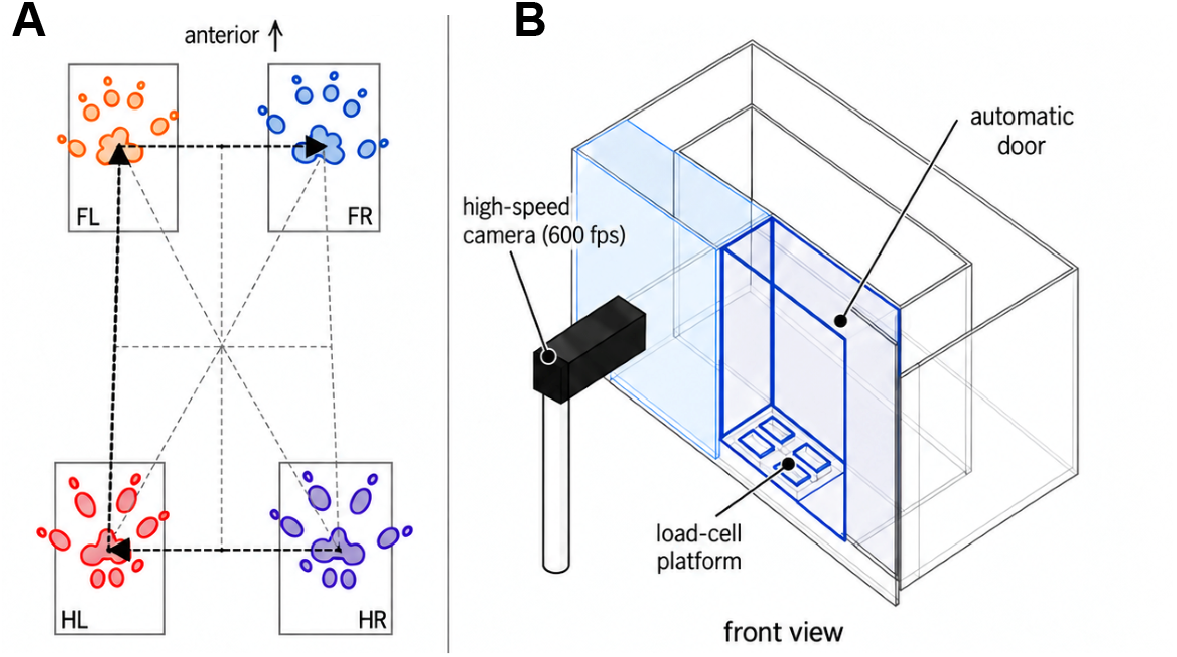
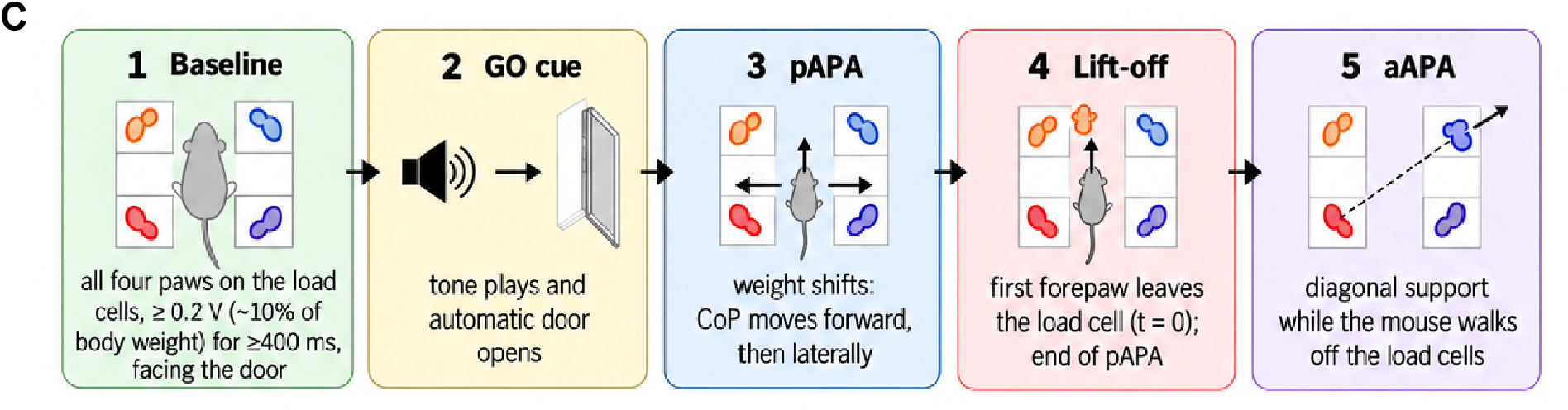
A load-cell paradigm for measuring APAs in freely moving mice. **(A)** Layout of the four load cells (FL, FR, HL, HR: front/hind, left/right). **(B)** Schematic front view of the arena. The mouse stands on the load cells behind an automatic door; after it exits through the door it can walk around a return walkway, and a manually operated door regulates its path. The arena is built from transparent acrylic and rests on posts that never touch the load cells. **(C)** Trial logic. A GO cue and door opening follow a ≥400 ms baseline stance. Front-paw lift-off (t = 0) defines the end of the pAPA.

Each load cell carries a small 3D-printed plate on which a single paw rests; the plates are separated by open space, which compels the mouse to place one paw per sensor, and the surrounding walkway is mechanically isolated from the load cells so that only the paws load them. The paradigm was developed in two phases. In the first, a touch sensor at snout height opened a door in front of the load cells. Mice leaned on the door and triggered it inadvertently, so we replaced the sensor with a pressure criterion: the door opens and an auditory GO cue plays only after the mouse has loaded the cells with ≥0.2 V (≈10% of body weight) for at least 400 ms (Fig. 1C). This required no learning and yielded a homogeneous baseline stance. Initial trial yield was low (16.7%) in a one-directional corridor. Adding an adjoining walkway that allows the mouse to return to the load cells, a second, manually operated door to regulate the mouse’s path, and a small plate between the load cells (to reduce the fear of falling into the gap) increased trial acquisition (Fig. 1B; Methods). The front wall is transparent, allowing a side-view high-speed camera (Basler, 600 fps in a reduced region of interest) to record the front paws, the snout and the door. A trial was analyzable if the mouse faced the door, stood still on all four load cells at threshold for ≥400 ms, did not dip its head between the load cells, and began locomotion by lifting a front paw before the other paws.

### Mice initiate steps with a diagonal weight-redistribution pattern

We aligned pressure traces from analyzable trials to the moment of front-paw lift-off (t = 0), which defines the end of the pAPA, and mirrored left-initiated trials so that the swing paw is always the front right paw (FR) (Fig. 2; Methods). The traces showed a consistent sequence. Pressure on the stance forepaw (FL) increases as weight is transferred onto it, pressure on the swing forepaw (FR) falls sharply at lift-off, the diagonally opposite hindpaw (HL) unloads shortly afterwards, and the mouse is then supported by the remaining diagonal pair (FL and HR). The sum of the four paws remained at approximately 100% of body weight until the mouse stepped off the load cells, as expected for a balanced redistribution. In the neutral stance mice also placed more of their weight on the hindpaws than on the forepaws. This diagonal pattern is the one described for cats and dogs (Gahery et al., 1980; Massion & Gahery, 1979).

**Figure 2.**
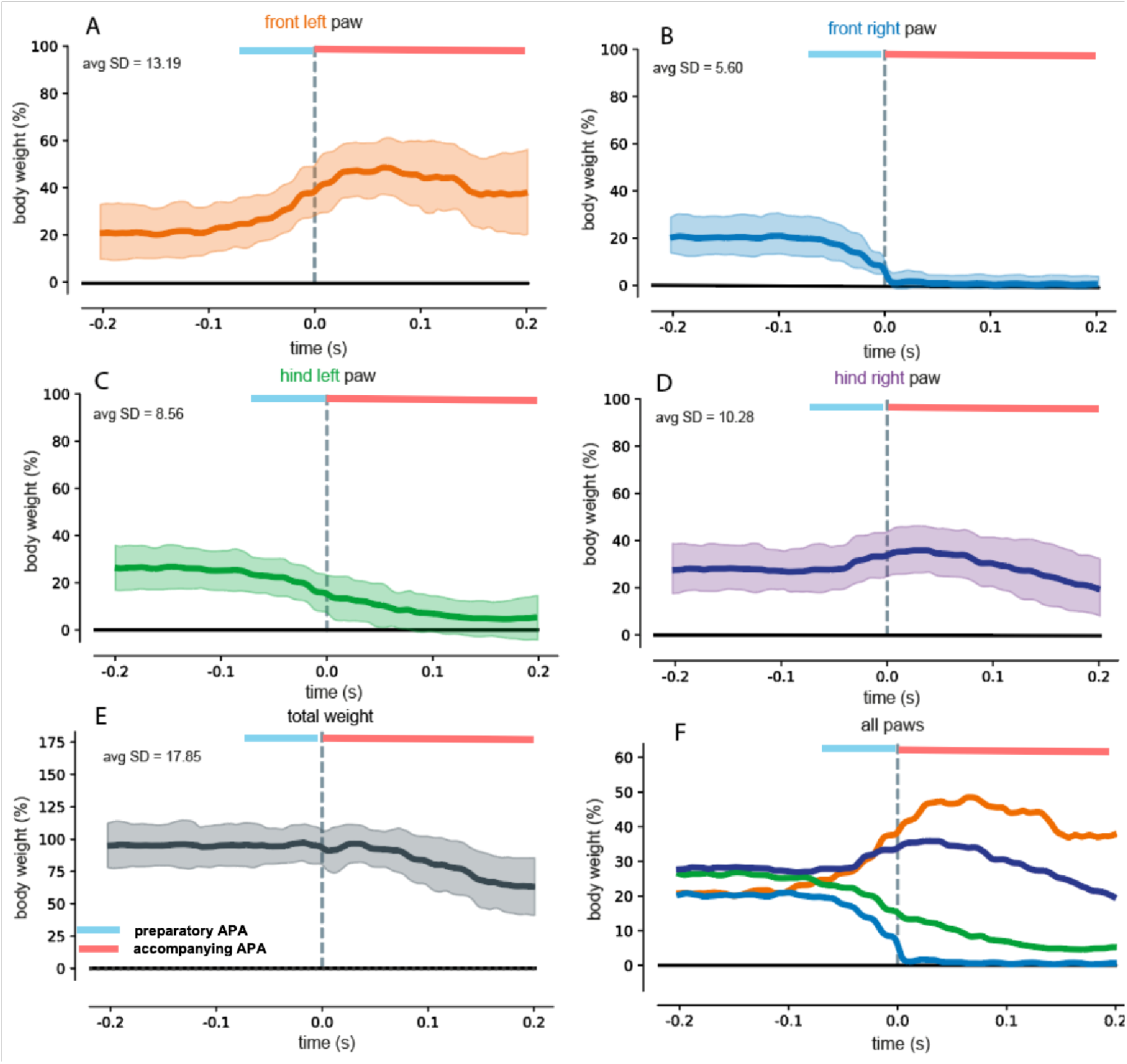
Paw-by-paw pressure during step initiation. Average pressure trace (bold) ± s.d. (shaded) of each paw, aligned to front-paw lift-off (t = 0), over 400 ms. Pale blue and pink bars mark the pAPA (blue) and aAPA (pink) periods. **(A)** FL (stance forepaw); **(B)** FR (swing forepaw); **(C)** HL; **(D)** HR; **(E)** sum of the four paws (% body weight); **(F)** all four paws together.

### The center of pressure shifts forward and then laterally

To summarize the redistribution of load across the four paws, we calculated the two-dimensional center of pressure (CoP) from the calibrated outputs of the four load cells. The voltage recorded by each load cell was baseline-corrected and converted to an equivalent load using a load-cell-specific calibration curve. When calibration values were expressed as mass in grams, they were converted to vertical ground-reaction force in newtons using:

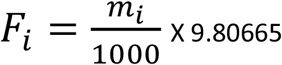

where F_i_ is the vertical force measured by load cell i in newtons, and m_i_ is the corresponding calibrated mass in grams. Each load cell was assigned a fixed spatial coordinate (x_i_, y_i_), corresponding to its physical position beneath the animal. At each time point, the center of pressure (CoP) was calculated as the force-weighted average of the fixed positions of the four load cells: front-left (FL), front-right (FR), hind-left (HL) and hind-right (HR). Each sensor’s spatial coordinates were multiplied by its measured vertical force, and these contributions were summed and divided by the total vertical force across all four sensors. The resulting x-coordinate described the medio-lateral position of the CoP, and the y-coordinate described its anterior–posterior position.

Thus, a load cell supporting a greater proportion of the total vertical force contributed more strongly to the estimated CoP position. If the load-cell coordinates were expressed in centimeters, the resulting CoP coordinates were also expressed in centimeters. Converting the calibrated loads from grams to newtons did not alter the estimated CoP position because the same conversion factor was present in both the numerator and denominator. Following the GO cue, the CoP initially shifted anteriorly, indicating a redistribution of vertical load toward the forepaws, and subsequently shifted laterally toward the left stance side. This lateral shift was associated with unloading of the right side and preceded lift-off of the front-right swing paw. As described for human gait initiation (Mizusawa et al., 2017), the trajectory ended with unloading of the impending swing limb. However, direct comparison between species should be made cautiously because mice use quadrupedal support, the direction and mechanical function of the initial CoP displacement may differ from those in bipeds, and the mouse trajectory occurs over a much smaller spatial scale. The load-cell measurements quantify the redistribution of vertical force but do not, by themselves, identify the individual muscles or horizontal forces responsible for producing the movement (Fig. 3; Methods).

**Figure 3.**
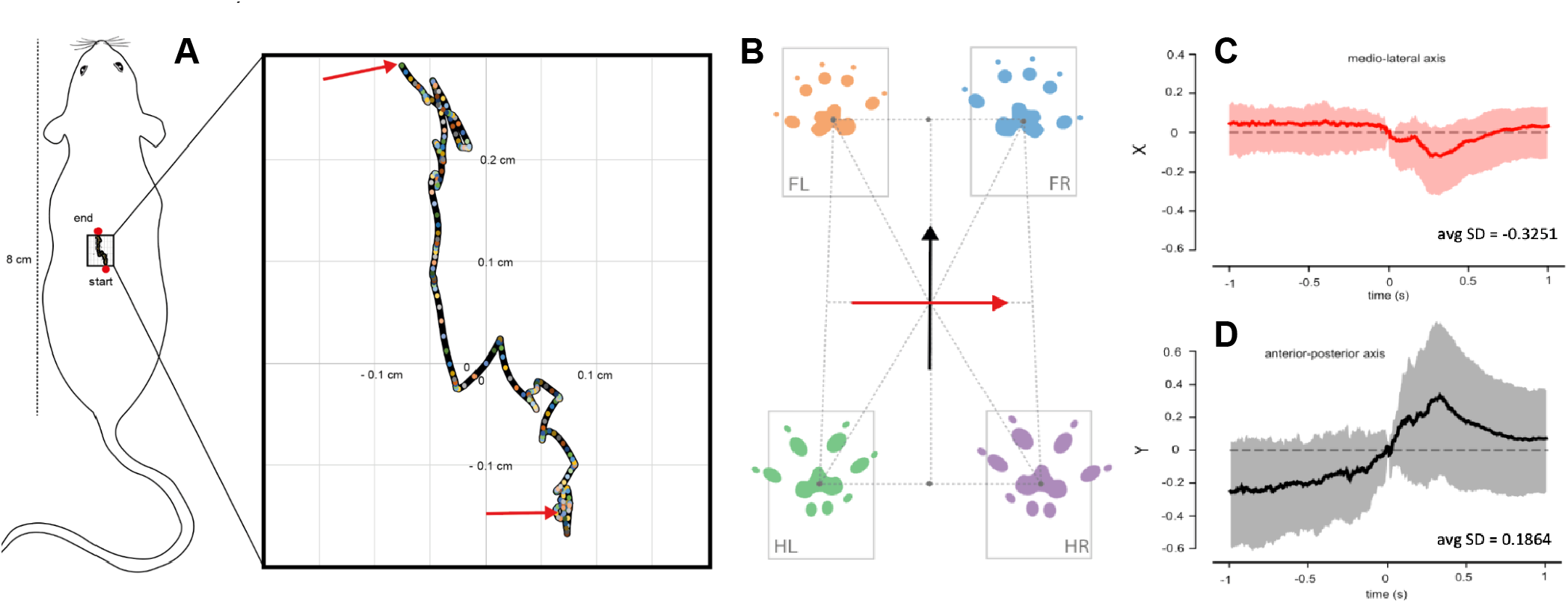
center-of-pressure dynamics during movement initiation. **(A)** Representative CoP trajectory plotted in two-dimensional *x–y* space. The mouse schematic indicates the location and scale of the analyzed region, which is enlarged on the right. Colored points show sampled CoP positions across trials, and the red arrows indicate the beginning and end of the trajectory. Together, these measurements show the redistribution of postural load that accompanies the transition from stable stance to forward locomotion. **(B)** Trial-wise paw-pressure distributions for the FL, FR, HL and HR paws, together with the coordinate system used to calculate whole-body CoP displacement. The red and black arrows indicate the mediolateral (*x*) and anterior-posterior (*y*) axes, respectively. **(C, D)** CoP displacement along the mediolateral and anterior–posterior axes, aligned to front-paw lift-off (*t* = 0). Solid lines represent the average trajectory across trials, shaded regions show across-trial variability and horizontal dashed lines indicate zero displacement.

### APAs are stereotyped across trials and have a pressure-defined pAPA of ∼88 ms

Individual load-cell traces contained measurement noise and trial-to-trial variability. To determine whether the underlying pattern of weight redistribution was nevertheless stereotyped, we applied principal component analysis (PCA) to the four-paw pressure traces from 60 analyzable trials. Each trial comprised 240 frames spanning the pAPA and aAPA periods. PCA reduced the four simultaneous paw-pressure signals to a single dominant component. The first principal component (PC1) explained approximately 80% of the variance in the average trial. By comparison, when the paw identities were shuffled within each trial and the same PC1 was applied to the shuffled data, the explained variance decreased to approximately 50% (Fig. 4A). This reduction indicates that the paw-specific sequence of loading and unloading contributes substantially to the stereotyped structure of the behavior. To estimate pAPA duration from the pressure signal, a continuous piecewise-linear function comprising four nodes and three segments was fitted to the front-right paw-pressure trace from each trial. The middle segment captured the rapid unloading of the front-right paw. Pressure-defined pAPA duration was therefore calculated as: 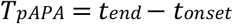 Here, t_end and t_onset are the fitted breakpoints delimiting the middle unloading segment. The mean pressure-defined pAPA duration was 87.9 ms, with a median of 76.6 ms and a range of 6.3–208.3 ms (Fig. 4B).

**Figure 4.**
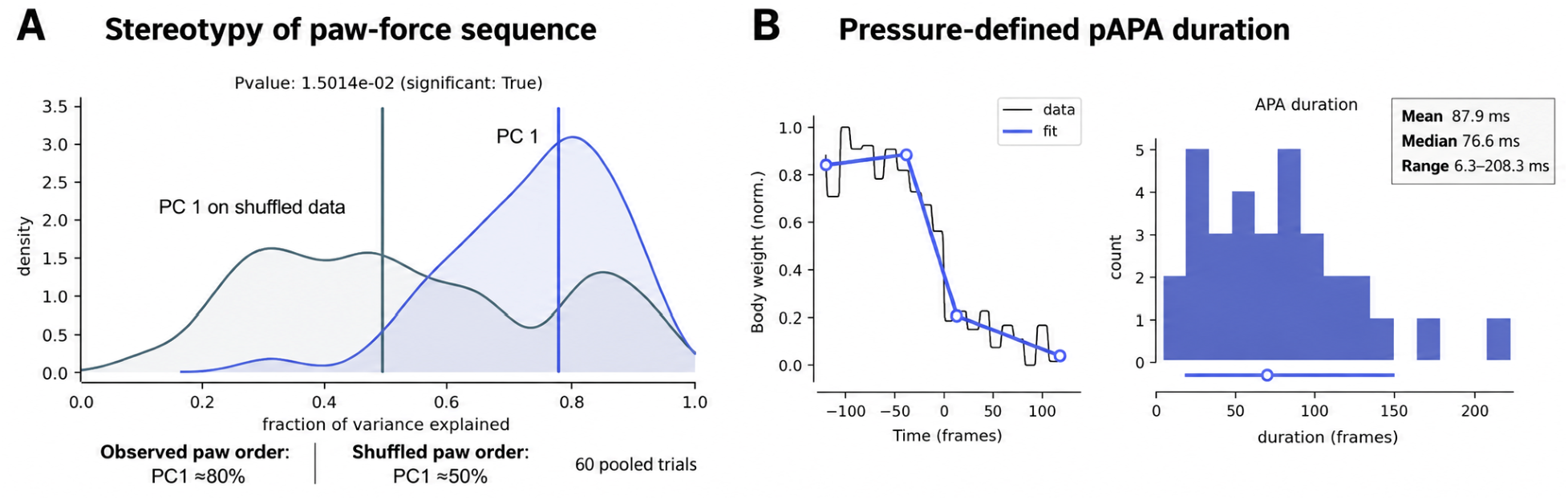
Mouse APAs are stereotyped. **(A)** Variance explained by the first principal component (PC1) of four-paw pressure traces (average trial of 60 trials), and by the same PC1 fitted to data in which paw order was shuffled (control). **(C)** A piecewise-linear fit to initiating forepaw pressure estimated pAPA duration. Data is pooled across trials.

### Markerless tracking reveals early movement before pressure-defined paw unloading

The pressure-based estimate captures only the component of the pAPA that changes the vertical forces applied through the paws. To identify earlier postural events, we tracked body landmarks in high-speed video using DeepLabCut (Mathis et al., 2018), including the eye, lower ear, and the toe, thumb and heel of the initiating forepaw (Fig. 5A). Detectable eye and ear displacement began approximately 200 ms before front-paw lift-off, substantially earlier than the pressure-defined pAPA. The anterior-posterior CoP shift became detectable approximately 120 ms before lift-off. The forepaw remained approximately flat, with an angle near 180°, until it began rotating away from this resting configuration approximately 40-22 ms before lift-off. It subsequently approached the near-vertical configuration associated with lift-off at (t=0). Using the onset of head/ear displacement as the beginning of the preparatory adjustment and the first detectable departure of the forepaw from its flat posture as the onset of the focal stepping movement, the preparatory interval was calculated as:

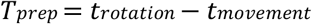

Substituting the observed event times gives:

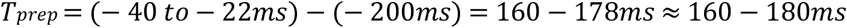

which was rounded to approximately 160–180 ms. The estimated 160–180 ms preparatory interval extends from the earliest detected head or ear movement to the onset of visible forepaw rotation. It excludes the final 22–40 ms during which the forepaw rotates toward lift-off. Head and ear displacement may also reflect orienting or other movements, so this interval provides an operational estimate of preparation rather than a definitive measure of pAPA duration. Although numerically similar to the approximately 150–200 ms durations reported in humans and values reported in other species, direct comparisons require caution because the events used to define preparation may differ. If the interval instead extended from the earliest head or ear movement to complete forepaw lift-off, its duration would be approximately 200 ms.

**Figure 5.**
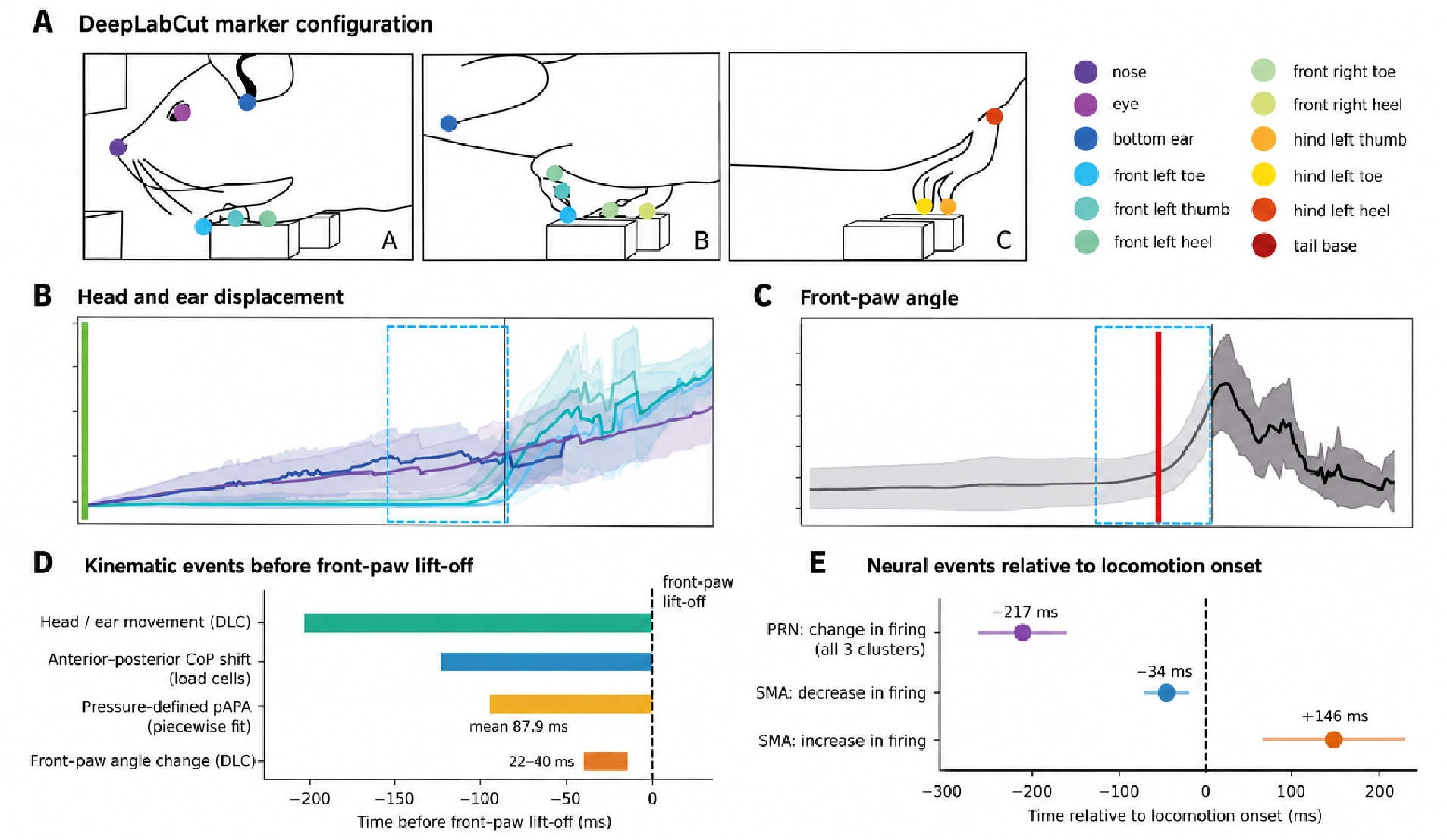
DeepLabCut-derived kinematics and timing of movement-related events. **(A)** DeepLabCut marker configuration used to track head, ear, forepaw, hind-paw and tail movements from side-view recordings. **(B)** Displacement trajectories of the tracked head/ear and forepaw markers relative to front-paw lift-off (*t* = 0). Lines represent the averaged trajectories and shaded regions show across-trial variability; the blue dashed box indicates the pressure-defined pAPA interval. **(C)** Front-paw angle trajectory relative to lift-off. The red vertical line marks the DLC-derived onset of the forepaw angle change, and the blue dashed box shows the pressure-defined pAPA interval for comparison. **(D)** Summary of kinematic-event timing relative to front-paw lift-off: head/ear movement (∼−200 ms), anterior–posterior center-of-pressure shift (∼−120 ms), pressure-defined pAPA onset (mean −87.9 ms) and forepaw angle change (−40 to −22 ms). Bars extend from the estimated onset of each event to lift-off, except for the forepaw-angle bar, which indicates the estimated onset range. **(E)** Neural-event timing relative to locomotion onset defined from body speed: mean change in pontine reticular nucleus (PRN) firing (−217 ms), supplementary motor area (SMA) firing decrease (−34 ms) and SMA firing increase (+146 ms). Horizontal lines indicate the range of individual cluster means. The temporal reference in **(E)** differs from that used in **(B-D)** body-speed-defined locomotion onset versus front-paw lift-off, and the neural recordings were obtained in a different arena; therefore, the zero points and event latencies should not be compared directly. Kinematic data are pooled across trials.

### PRN neurons change their firing before locomotion onset

The PRN contains reticulospinal neurons and has been implicated in postural control. To test whether PRN activity is related to the APA time window, we analyzed Neuropixels recordings from freely moving mice walking along a linear track (recorded by F. Claudi for a separate study; Methods). Locomotion onset was defined from body speed (<1 cm/s at rest and ≥2.5 cm/s for movement), keeping only transitions with a quiet period beforehand. We recorded 304 PRN neurons from 4 mice over 614 transitions (256 left-paw and 358 right-paw initiations) and clustered the time-averaged peri-stimulus time histograms (PSTHs), which produced five clusters (95, 62, 57, 43 and 47 neurons). Three of the five clusters showed a significant change in firing rate between baseline and the post-onset comparison period, comprising 214 of 304 neurons (70%; Fig. 6). The mean deviation from baseline began 217 ms before locomotion onset (191, 264 and 197 ms for the three clusters), which is a similar time scale to the head and ear movement that marks the beginning of the pAPA (Fig. 5B). We performed the same analysis for the SMA (376 neurons, 1 mouse, 276 transitions). Four of five clusters changed their firing around locomotion onset: 167 of 376 neurons decreased their activity and 128 increased it, i.e. 295 of 376 (78%) were modulated. The decreases began on average 34 ms before locomotion onset (48 and 20 ms for the two clusters), and the increases began 146 ms after onset (80 and 212 ms), i.e. during the aAPA phase.

**Figure 6.**
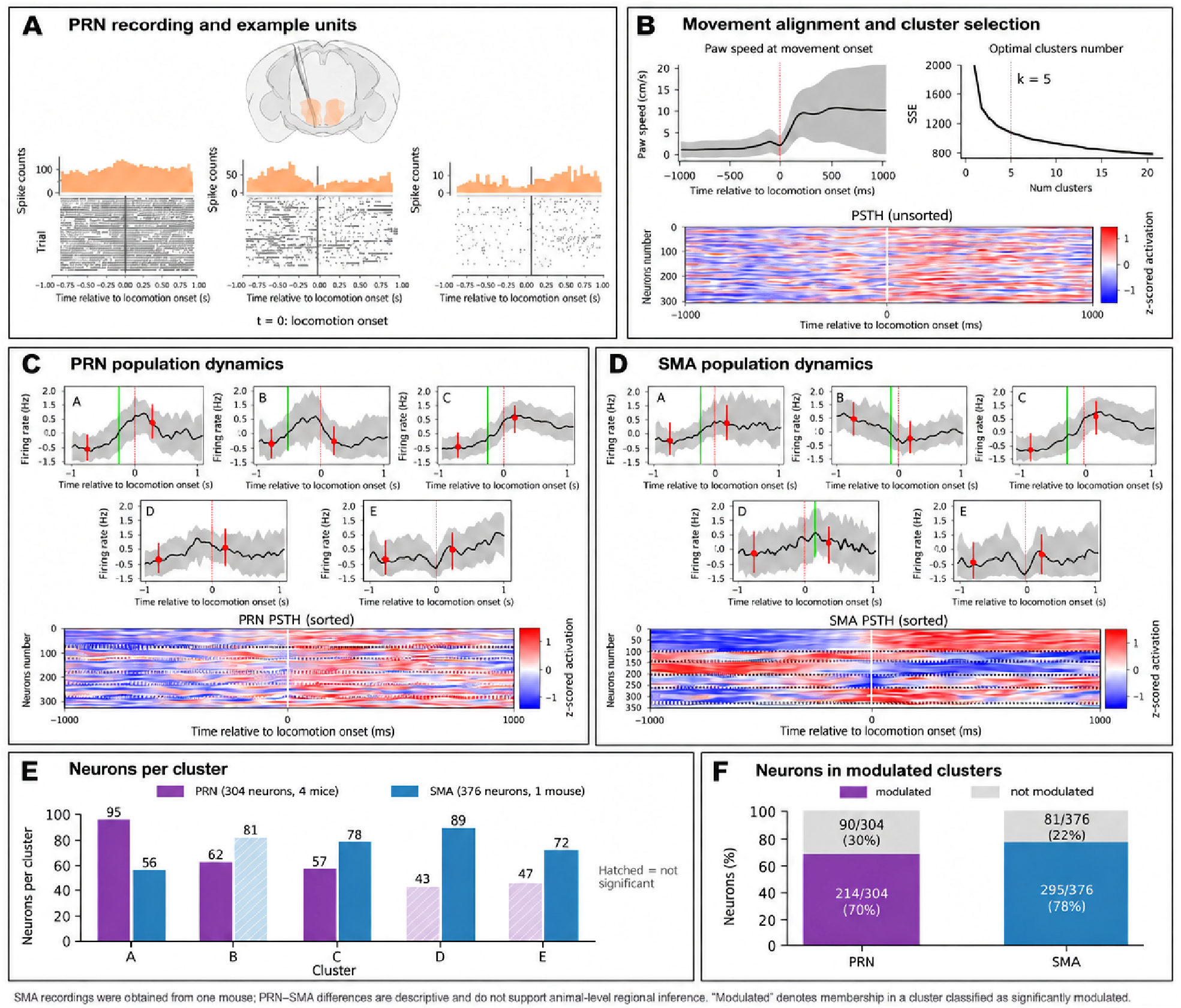
Peri-locomotion neural dynamics in the pontine reticular nucleus and supplementary motor area. **(A)**Coronal brain section showing the pontine reticular nucleus (PRN) recording location and reconstructed probe trajectories, together with spike-count histograms and raster plots from three representative units. Neural activity is aligned to locomotion onset (*t* = 0). **(B)** Definition of locomotion onset from body speed, selection of a five-cluster solution from the sum-of-squared-error curve and unsorted peri-locomotion population activity. **(C, D)** Cluster-averaged firing-rate trajectories and sorted population activity for PRN and supplementary motor area (SMA), respectively. Black lines show the mean z-scored firing rate within each cluster; grey shading indicates the standard deviation across neurons. The red dotted line marks locomotion onset, defined from body speed. Red points and error bars indicate the mean and standard deviation of z-scored activity during the baseline and post-onset comparison periods. Green vertical lines mark the estimated time of activity change. Heatmaps show peri-locomotion *z*-scored activity, with red indicating greater and blue indicating lower activity. **(E)** Number of neurons assigned to each cluster. Hatched bars identify clusters that were not significantly modulated; significantly modulated clusters were PRN A-C and SMA A, C, D and E. Cluster labels are region-specific and do not imply equivalent response classes across PRN and SMA. **(F)**Proportion of neurons assigned to significantly modulated clusters: 214 of 304 PRN neurons (70%) and 295 of 376 SMA neurons (78%). PRN recordings comprised 304 neurons from four mice, whereas SMA recordings comprised 376 neurons from one mouse.

### Reticulospinal PRN neurons receive input from more than 70 areas

To identify the inputs to the pontine reticulospinal neurons (RSNs), we used monosynaptic rabies tracing restricted to the RSN population. A retrograde AAV-Cre was injected into the lumbar spinal cord (L3) to label RSNs projecting there, a Cre-dependent TVA/glycoprotein helper virus was injected into the PRN, and EnvA-pseudotyped rabies virus (tdTomato) was subsequently injected into the PRN (Methods). Across four mice we counted 2,065 labelled input neurons in more than 70 brain areas. The largest numbers of labelled neurons were within the PRN itself (387; 19%), followed by the periaqueductal gray (259; 13%), gigantocellular reticular nucleus (207; 10%), midbrain reticular nucleus (182; 9%) and intermediate reticular nucleus (130; 6%) (Fig. 7). Cortical input included the secondary motor area (58 neurons; ∼3%) and primary motor area (30; ∼1.5%). Other inputs came from the zona incerta, superior colliculus, vestibular nuclei, red nucleus, parabrachial nucleus and cuneiform nucleus. The distribution was variable between animals (84– 923 labelled cells per brain), so the percentages should be taken as approximate.

**Figure 7.**
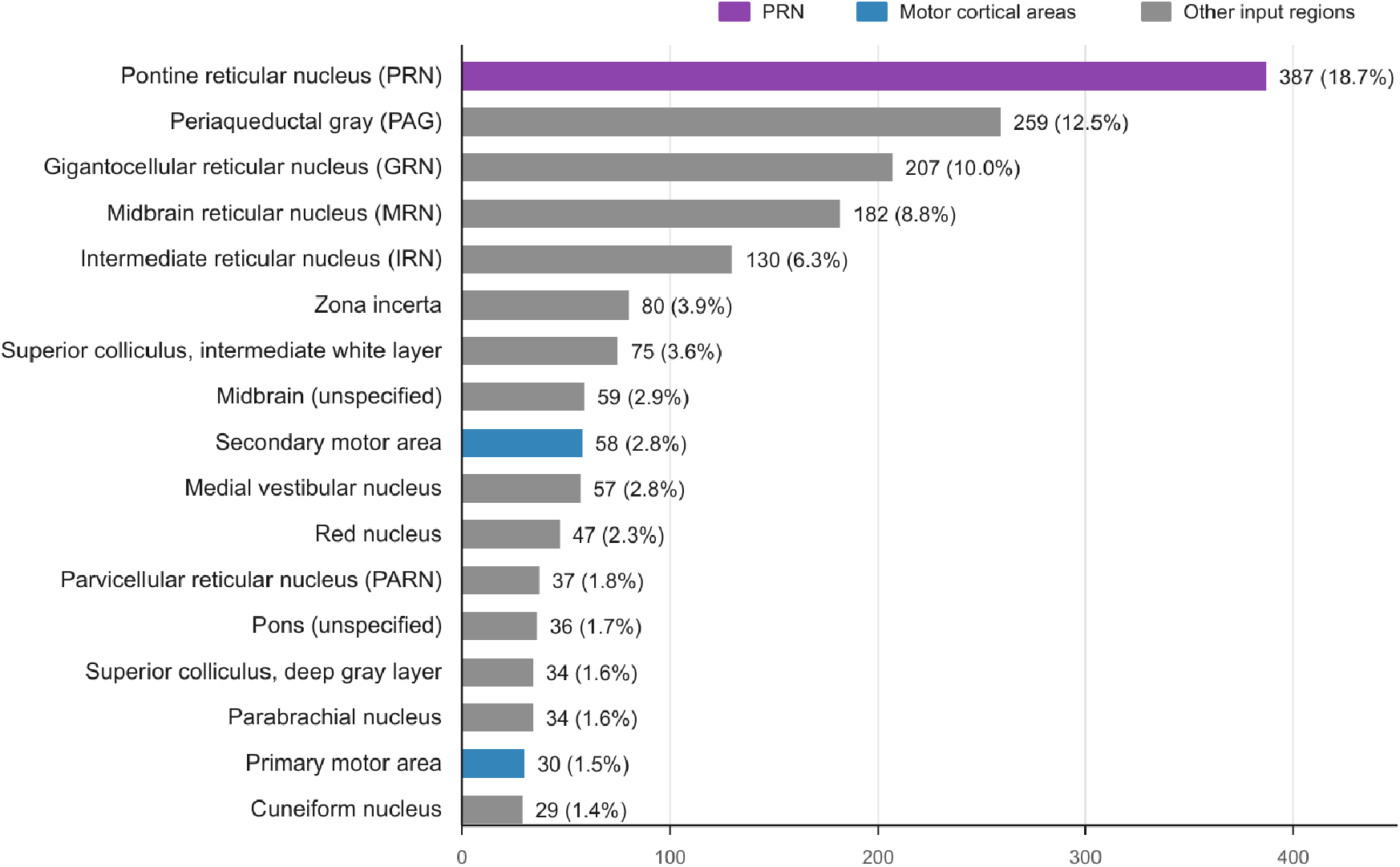
Major anatomical inputs to reticulospinal pontine reticular nucleus neurons. Horizontal bars show the numbers of rabies-labelled input neurons detected in the 17 most highly represented regions among more than 70 identified input areas. Counts were pooled across four mice and are presented together with their percentage of the total labelled population (*n* = 2,065 cells). PRN is highlighted in purple, motor cortical inputs are highlighted in blue and include the secondary motor area (58 cells, 2.8%) and primary motor area (30 cells, 1.5%); all other regions are shown in grey. Percentages represent the pooled cellular distribution and do not capture between-animal variability.

### Preliminary ablation of PRN neurons alters paw coordination

To explore the contribution of PRN neurons to APAs, we used diphtheria toxin receptor (DTR)-mediated cell ablation with two targeting strategies. Broad neuronal targeting used AAV-CMV-Cre together with AAV-FLEX-DTR-EGFP in wild-type mice (DTR-Pan), whereas glutamatergic-neuron targeting used AAV-FLEX-DTR-EGFP in Vglut2-Cre mice (DTR-Glut). Following baseline behavioral testing, diphtheria toxin (DT) was administered subcutaneously (∼110 μg/kg). Representative histological sections showed a marked reduction in labelled cells after DT administration in both targeting conditions (Fig. 9). Control sections contained approximately 92 and 55 labelled cells on average for the DTR-Pan and DTR-Glut experiments, respectively, whereas few labelled cells were visible in the corresponding post-treatment sections. These observations support depletion of the labelled populations but do not establish the completeness or spatial specificity of ablation. Both lesion groups still produced a stereotyped sequence of weight redistribution when analyzed with PC1 from control mice (PC1 explained ∼75% of the variance in DTR-Pan and ∼72% in DTR-Glut, versus ∼50% for shuffled data; Fig. 8A), although the DTR-Glut CoP trajectory appeared less smooth. The pressure-defined pAPA was numerically shorter in the DTR-Pan and DTR-Glut groups (mean 80.4 and 65.3 ms; Fig. 8B). To quantify interlimb coordination, we computed the Pearson correlation between the FR and HL pressure traces in each trial. The FR–HL correlation distribution of DTR-Glut mice differed significantly from that of controls (t-test), whereas the DTR-Pan group did not (Fig. 9). These results hint that glutamatergic PRN neurons contribute to the timing of the diagonal FR–HL sequence at the start of the step, but they come from only two mice and 15 trials per condition and must be treated as preliminary (see Discussion).

**Figure 8.**
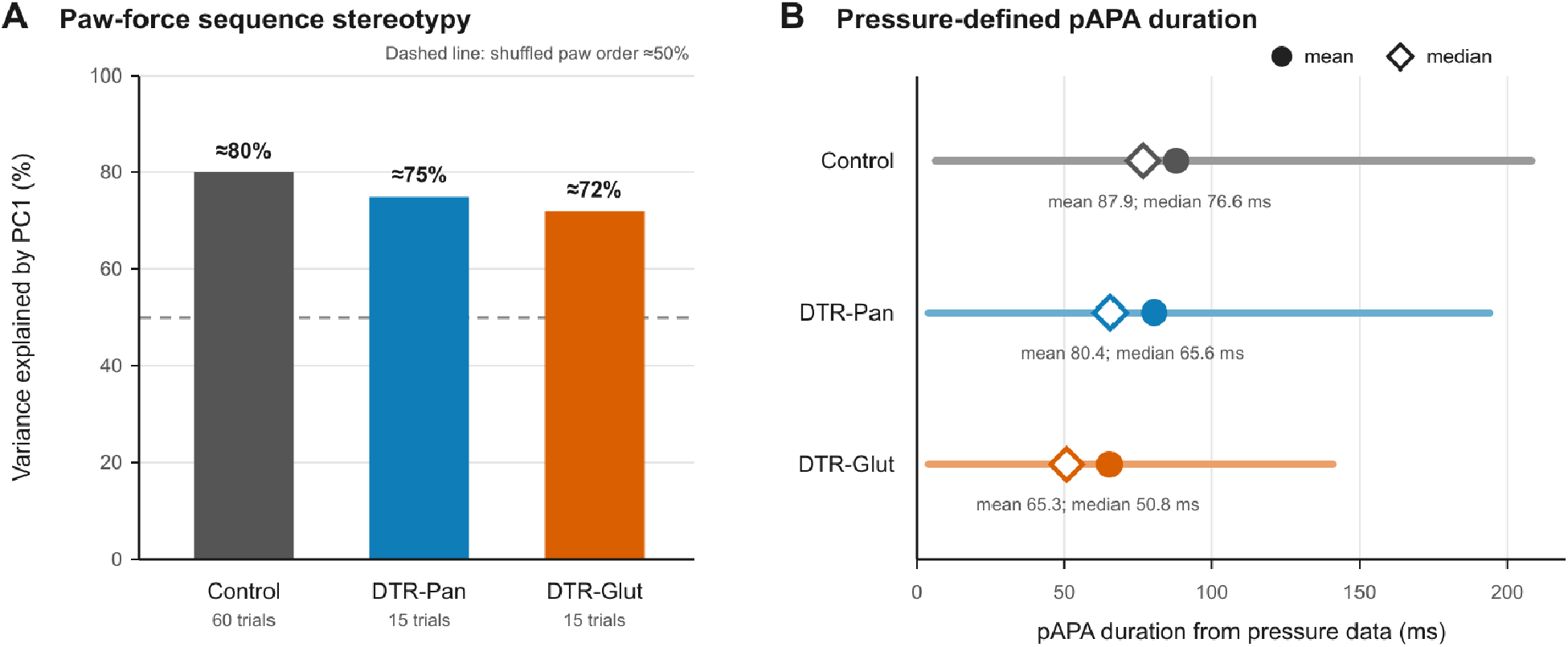
Paw-force stereotypy and pressure-defined anticipatory postural adjustment timing after PRN ablation. **(A)** Variance in the four-paw pressure sequence explained by the first principal component (PC1) in control, pan-neuronal DTR-ablation (DTR-Pan) and glutamatergic-neuron DTR-ablation (DTR-Glut) conditions. The dashed horizontal line indicates the approximately 50% variance explained after shuffling paw order. The analysis included 60 control trials and 15 trials from each ablation condition; the DTR-Pan and DTR-Glut datasets comprised two mice per group. **(B)** Duration of the pressure-defined anticipatory postural adjustment (pAPA), estimated by piecewise-linear fitting of the initiating-forepaw pressure trace. Horizontal lines show the observed range, filled circles indicate the mean and open diamonds indicate the median.

**Figure 9.**
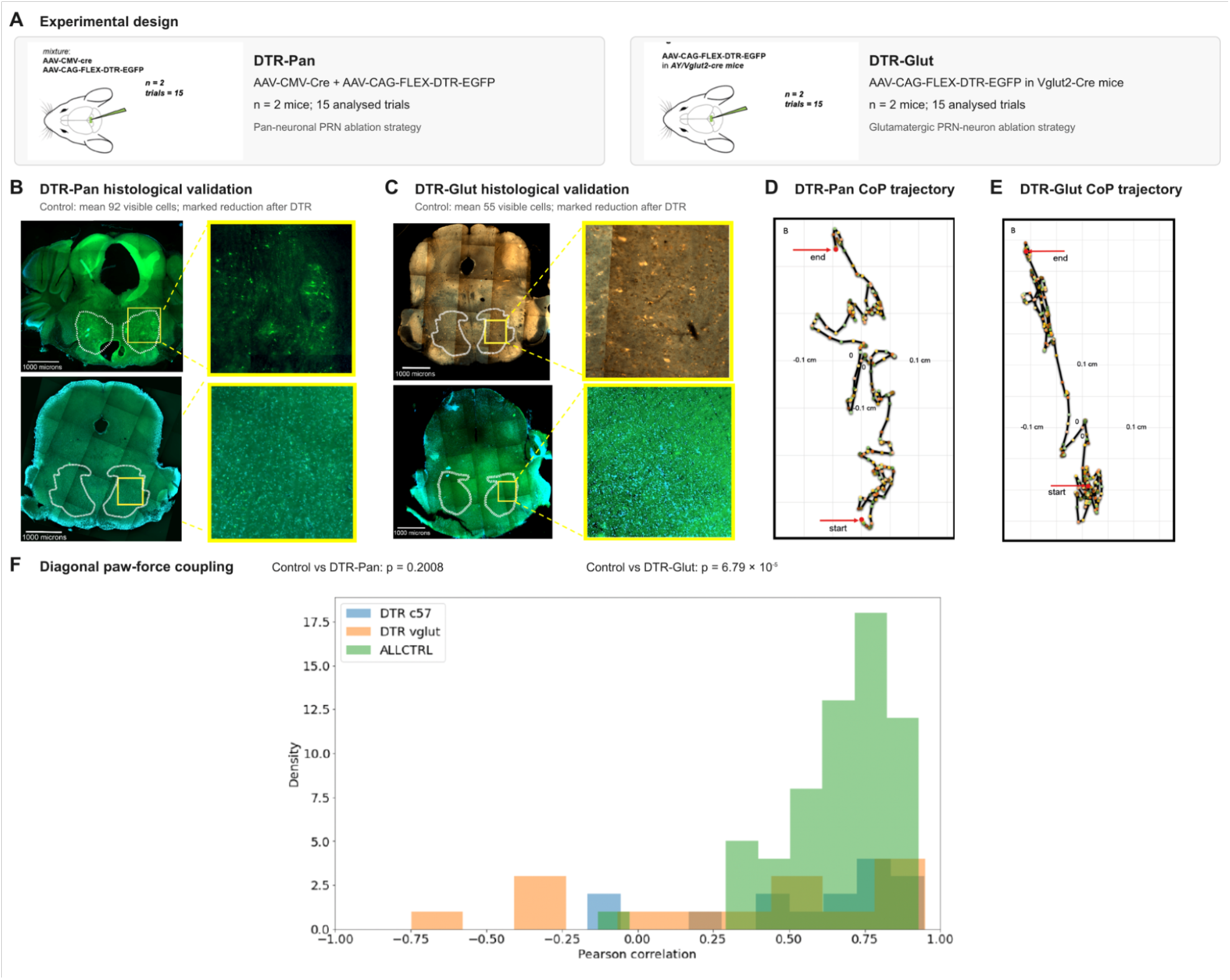
PRN ablation strategies, histological validation and effects on postural coordination. **(A)** Experimental designs for pan-neuronal and glutamatergic-neuron ablation in the pontine reticular nucleus (PRN). **(B)** Representative coronal sections and magnified PRN regions for the DTR-Pan experiment. The upper images show the control condition and the lower images show the marked reduction in labelled cells following DTR treatment. **(C)** Corresponding histological validation of the DTR-Glut experiment. The upper images show the control condition and the lower images show the reduction following diphtheria toxin (DT) administration. Scale bars in the coronal sections represent 1,000 μm. **(D, E)**Representative center-of-pressure (CoP) trajectories during movement initiation in the DTR-Pan and DTR-Glut conditions, respectively. Sampled CoP positions are displayed in the mediolateral and anterior– posterior plane, with successive samples connected to illustrate the progression of the trajectory; arrows identify its start and end. **(F)**Distributions of Pearson correlation coefficients between the front-right and hind-left paw-force traces across control, DTR-Pan and DTR-Glut trials.

**Table 1.** Experimental cohorts and datasets.

| Dataset | Animals | Sample | Used for |
|---|---|---|---|
| Force-plate paradigm, controls | C57BL/6J, 8–16 wk, female and male; n=91 | 60 analyzable trials (PCA); trials aligned to front-paw lift-off | Figs 1–5 |
| Neuropixels, PRN | 4 mice (freely moving, linear track) | 304 neurons; 614 locomotion-onset trials (256 left, 358 right paw) | Fig. 6 |
| Neuropixels, SMA | 1 mouse | 376 neurons; 276 trials (123 left, 153 right paw) | Fig. 6 |
| Rabies tracing from RSNs | 4 male C57BL/6J, 6–8 wk | 2,065 labelled input neurons; >70 areas | Fig. 7 |
| DTR-Pan ablation | 2 female C57BL/6J | 15 analyzable trials | Figs 8, 9 |
| DTR-Glut ablation | 2 male Vglut2-Cre | 15 analyzable trials | Figs 8, 9 |

## Discussion

We developed a simple force-plate paradigm in which mice stand on four independent load cells and then step off voluntarily. The paradigm resolves the weight carried by each paw during a step and is compatible with high-speed video and neural recordings. With it we showed that mice initiate a step with a stereotyped, diagonal pattern of weight redistribution: the CoP moves forward and then laterally away from the swing side, the swing forepaw and diagonal hindpaw unload in sequence, and the remaining diagonal pair supports the body. This is the same strategy that has been described for cats and dogs (Gahery et al., 1980; Massion & Gahery, 1979), and, as in human gait initiation (Mizusawa et al., 2017), the CoP follows a stereotyped trajectory that unloads the swing limb. That a species with a different body plan and posture uses a comparable solution supports the idea that a shared set of principles underlies APAs and justifies using mice as a model.

A methodological point of the study is that APA timing depends on the measurement used. The pressure signal, which is what has typically been used in force-plate studies, places the end of the pAPA at lift-off and the beginning ∼88 ms earlier. Pose tracking showed movement of the head and ear approximately 200 ms before lift-off, well before the load cells registered a change. The load cells measure vertical paw loading and may therefore miss early changes in body configuration that produce little detectable redistribution of support. Head and ear movement preceded the prominent unloading of the initiating forepaw. The interval from the earliest detected head movement to the onset of forepaw rotation was approximately 160–180 ms. Because head movement may also reflect orienting, this interval is an operational estimate of preparation rather than a definitive pAPA duration. Its numerical similarity to durations reported in humans should be interpreted cautiously because the events used to define the interval differ between studies (Mizusawa et al., 2017). These observations argue for measuring APAs with several complementary methods; electromyography (EMG), which we did not record, would add further information about the underlying muscle activity, in particular of axial muscles.

The neural recordings identified movement-related activity in PRN and SMA. In the PRN, 214 of 304 neurons belonged to clusters classified as significantly modulated, with an average estimated activity-change time approximately 217 ms before locomotion onset. SMA clusters showed both pre-onset decreases and post-onset increases in activity. These recordings were obtained during a separate locomotion task without simultaneous paw-force measurements, so they identify candidate neural populations for further investigation but do not establish that the recorded activity specifically represents APAs. The rabies tracing showed that reticulospinal PRN neurons receive converging input from a large number of areas, including motor cortex, midbrain, cerebellar-related and other reticular regions, consistent with the proposal that the PRN integrates multiple signals and relays them to spinal motor circuits (Brownstone & Chopek, 2018; Sivertsen et al., 2014). The preliminary ablation experiments suggested altered FR–HL force coupling in the DTR-Glut condition, whereas no statistically significant difference from control was detected in the DTR-Pan condition. This pattern motivates further investigation of glutamatergic PRN populations, but it does not establish that the two targeting strategies have different effects. Each manipulation group contained only two mice, and the statistical comparisons pooled trials. Larger cohorts, animal-level analyses and quantitative validation of the extent and specificity of ablation are therefore needed to determine whether PRN populations contribute causally to this coordination.

### Limitations

Several limitations should be kept in mind. First, the ablation experiments are underpowered: each lesion group comprises two mice and 15 trials, trials are not independent samples from independent animals, and the statistical comparison of FR–HL correlation was done across trials. We therefore consider them preliminary and hypothesis-generating. Second, the neural recordings were obtained for a different study, in a different arena (a linear track, 60 fps video) and with locomotion onset defined from body speed rather than paw lift-off; they were not recorded during the force-plate paradigm, and so the neural and kinematic timing can be compared only approximately. The SMA data come from a single mouse. Third, pressure recordings are sensitive to the exact position of the paws on the plates, and the paradigm yields relatively few analyzable trials per session. Fourth, left- and right-initiated trials were pooled by mirroring, which assumes symmetry. Finally, the modest sample of trials means that the quantitative estimates (e.g. pAPA duration, which varied from <10 to >200 ms) should be interpreted as descriptive.

### Outlook

The force-plate platform is inexpensive, built primarily from commercially available components and compatible with the experimental tools available in mice, including cell-type-specific viral manipulation, optogenetic and chemogenetic perturbation, calcium imaging, electrophysiology and electromyography. These combinations should make it possible to test whether activity in defined PRN populations is necessary for the temporal and spatial organization of anticipatory postural adjustments and to distinguish preparatory activity from signals related to locomotor execution. Postural instability is a major contributor to falls, morbidity and reduced quality of life in Parkinson’s disease (Park et al., 2015). The PRN receives input from brain regions that have been investigated as deep-brain-stimulation targets, including the pedunculopontine nucleus and caudal zona incerta, although the clinical evidence for effects on gait and balance remains variable (CollombClerc & Welter, 2015; Plaha et al., 2006). Applying the present paradigm to established dopaminedepletion models could determine whether Parkinsonian pathology alters the timing, magnitude or paw-specific organization of APAs and whether manipulation of PRN circuits modifies these deficits. Such experiments should use validated lesion models and include quantitative histological confirmation of dopamine depletion. A separate moving-walkway paradigm analogous to the “broken escalator” experiment (Bronstein et al., 2009) was also developed to study contextual adaptation of APAs, but those experiments were not completed.

## Methods

### Animals

All experiments were performed under the UK Animals (Scientific Procedures) Act 1986 under project licences PPL 70/7652 and PFE9BCE39. Male and female adult C57BL/6J mice aged 8–16 weeks were used for the force-plate experiments. Mice were habituated to the arena for approximately one week, initially for 5 min per day with exposure to the auditory GO cue and automatic door. They were subsequently tested once per weekday for 10 min per session, extended to approximately 20 min when performance remained stable, until performance declined, typically after approximately two months. Rabies tracing was performed in four male C57BL/6J mice aged 6–8 weeks. Pan-neuronal ablation experiments used two female C57BL/6J mice. Glutamatergic-neuron ablation experiments used two male Vglut2-Cre mice aged 8–16 weeks. Four adult mice aged 14–16 weeks, comprising two C57BL/6J mice and two Vgat-Cre/Ai14 mice, were used to characterize spinally projecting PRN cell types.

### Behavioral apparatus

Four single-point aluminium load cells (Model 1004) were mounted on posts on a 750 × 750 mm aluminium breadboard. Each carried a 3D-printed plate (1.7 × 1.2 cm in the final version) on which one paw could rest; the plates were separated by open space and the distances between loadcell centers were 1.8 cm (FL-FR), 3.5 cm (FL-HL) and 2.0 cm (HL-HR). The arena, laser-cut from transparent 5 mm acrylic, was supported by separate posts at the height of the load cells, never touching them, and was secured against tipping. It consisted of a one-directional start corridor, an automatic door immediately in front of the load cells, an adjoining walkway that returns to the corridor, and a second manually operated door. A custom acrylic piece between the load cells provides a safe passage across the gap. The load-cell signals were conditioned by a 4-channel amplifier (MA 102, zeroed before each session) and acquired with a CED Power1401-3A at 10 V excitation. The automatic door was driven by a Dynamixel XL-320 servo controlled through an OpenCM 9.04 board, a speaker played the GO cue, and a Basler acA1920-150um camera was triggered from an Arduino and recorded the left side of the mouse through the transparent front wall at an average of 600 fps with a reduced region of interest (192 × 160 px; offsets x = 448, y = 640) covering the front paws, snout and door. A top-view camera used during development was removed because tracking showed no additional information and frame rate could then be increased from 200 to 600 fps. Air puffs were occasionally used to keep mice from standing for long periods. Data was stored on an external SSD. Experiments were controlled with Python scripts (main.py, forceplate_config.py), with Arduino firmware (StandardFirmata and custom scripts for the speaker, camera trigger and door); all code, together with the full parts list, is available at https://github.com/Egzona66/APA_setup.

### Trial triggering and selection

A trial was triggered when the weight on each of the four load cells exceeded 0.2 V (≈10% of body weight, determined as the minimum output while standing in 5 mice) for at least 400 ms, which opened the door and played the GO cue. Videos were inspected frame-by-frame (manual_video_inspection.py) and a trial was considered analyzable if the mouse (1) faced the door, (2) stood still with ≥0.2 V on each load cell for ≥400 ms, (3) did not dip its head into the gap between the load cells before initiating locomotion, and (4) initiated locomotion by lifting a front paw first. For each trial the video, the frame of lift-off, the direction of walking, the stepping paw, the mouse’s weight and the frame rate were recorded in a metadata table.

### Pressure analysis

Voltage outputs were converted to grams using a calibration in which known weights (1–35 g) were placed on each load cell and the voltage was recorded (Spike2, CED). Weights were expressed as a percentage of each mouse’s body weight. Data were resampled to a common frame rate, smoothed with a Gaussian kernel, and aligned to the frame of front-paw lift-off. Trials in which the mouse stepped with the left forepaw were mirrored (FL↔FR, HL↔HR) so that the swing paw was always FR. The CoP was computed from the calibrated weights *W* of the four load cells as an anterior-posterior coordinate and a medio-lateral coordinate and plotted over 400 ms.

### Principal component analysis

To test for stereotypy, PCA was applied to the four-dimensional vertical-force traces (240 frames spanning pAPA and aAPA) of 60 trials, reducing them to one dimension. The variance explained by PC1 for the average trial was compared with that obtained when the same PC1 was fitted to data in which the order of the paws was shuffled within each trial. PCA is a linear method, but one component accounted for ∼80% of the variance, which limits concern about non-linearity. Clustering was not used since the aim was to test for stereotypy rather than to identify a number of movement types. For lesion groups the PC1 obtained from control mice was used.

### pAPA duration

A piecewise-linear function with four nodes (three segments) was fitted to the FR pressure trace of each trial. The second segment, in which pressure falls steeply as the paw unloads, was taken as the pAPA, and its duration was computed for every trial (apa_60ms.ipynb). Piecewise fitting was preferred to differentiating the signal because derivatives of the noisy load-cell signals fluctuate strongly.

### Markerless pose estimation

Twelve body parts were labelled manually in a subset of video frames that covered the range of postures and used to train a DeepLabCut network (Mathis et al., 2018), which was then applied to the remaining videos. The parts analyzed here were the nose, eye, lower ear, and the toe, thumb and heel of the left forepaw. Movement onset of the eye and ear was taken as the time at which their position departed from baseline. The paw angle was computed from the toe, thumb and heel (180° when flat), and the time at which it began to change (derivative analysis) was used as the end of the pAPA.

### Neuropixels recordings and analysis

Neural recordings were acquired by F. Claudi during a separate study of goal-directed locomotion in freely moving mice (Claudi, 2022). The mice walked along a 260 × 8 cm linear track positioned within a transparent acrylic arena measuring 40 × 60 × 6 cm and elevated 60 cm above the floor. Experiments were conducted in darkness under infrared illumination, and behavior was recorded from beneath the arena at 60 frames/s. Neuropixels probes (Jun et al., 2017a) targeted to the cuneiform and pedunculopontine nuclei also traversed the pontine reticular nucleus (PRN); recordings obtained from the PRN were reanalyzed here together with recordings from the supplementary motor area (SMA). Electrophysiological data were acquired using SpikeGLX version 3. Behavioral and electrophysiological recordings were synchronized using a 1 Hz, 5 V square-wave TTL signal with a 50% duty cycle, which was recorded by both acquisition systems. Automatic spike detection and sorting, followed by manual curation, were performed in MATLAB using Janelia Rocket Cluster 3.0 (JRCLUST; Jun et al., 2017b). Only units satisfying an absolute refractory-period criterion of at least 1 ms were included. Unless otherwise stated, all units passing this criterion were retained for analysis. Video-derived behavioral data were upsampled from 60 Hz to 1 kHz. The snout, neck, body, tail base and four paws were tracked using a custom-trained DeepLabCut network (Mathis et al., 2018). Each unit’s firing rate was calculated at 1 kHz by convolving a binary spike-occurrence vector with a Gaussian kernel with a standard deviation of 100 ms. Stationary periods were defined as epochs in which body speed was <1 cm/s. Locomotion onset was defined from body speed using a movement threshold of ≥2.5 cm/s. Neural activity was analyzed from −2 to +2 s relative to locomotion onset. The PRN dataset comprised 304 neurons recorded from four mice across 614 locomotion onset trials, including 256 left-paw-initiated and 358 right-paw-initiated trials. The SMA dataset comprised 376 neurons recorded from one mouse across 276 trials, including 123 left-paw-initiated and 153 right-paw-initiated trials. Each neuron’s trial-averaged firing-rate trace was z-scored and grouped using time-series k-means clustering with Euclidean distance. Five clusters were selected by visual inspection of the clustering-error curve using the elbow method. Baseline and post-onset activity were compared for each cluster using an independent-samples t-test calculated from summary statistics. Bonferroni correction was applied across the five cluster comparisons, with a corrected significance threshold of p < 0.05. For clusters classified as modulated, the estimated time of activity change was the first point at which the cluster representative crossed a threshold one baseline standard deviation above or below the baseline mean, according to the direction of modulation.

### Retrograde monosynaptic tracing

Monosynaptic rabies tracing was performed in four male C57BL/6J mice aged 6-8 weeks to identify putative monosynaptic inputs to spinally projecting PRN neurons. AAV-Retro-Cre was first injected into the L3 spinal cord. In a subsequent surgery, a Cre-dependent helper virus AAV-Flex-TVA-hHA-GLY was injected into the PRN to express the avian TVA receptor and rabies glycoprotein selectively in spinally projecting PRN neurons. EnvA-pseudotyped RABVtdTomato was subsequently injected into the PRN to label starter neurons and their putative monosynaptic inputs (Wickersham et al., 2007). The PRN injection coordinates were Y = −4.5, X = ±1 and Z = −5. Following tissue collection, immunohistochemistry was performed and labeled brain sections were imaged using a Zeiss confocal microscope. Labeled cells were counted separately for each anatomical area and each mouse. Their anatomical locations were reconstructed in a three-dimensional reference brain using SHARP-Track (Shamash et al., 2018). Across the four mice, 2,065 labeled cells were assigned to more than 70 anatomical areas.

### Diphtheria toxin ablation

DTR-Pan mice were injected with a 1:1 mixture of AAV-CMV-Cre and AAV-CAG-FLEX-DTR-EGFP (Saito et al., 2001) into the PRN; DTR-Glut (Vglut2-Cre) mice were injected with AAV-CAGFLEX-DTR-EGFP (140 nL per side; Y −4.5, X ±1, Z −5). After two days of recovery, mice were habituated for half a week and tested for another 1.5 weeks before DT. DT was injected subcutaneously (stock 2 μg/μL diluted 120× to 16.67 ng/μL; 200 μL, i.e. 3.3 μg per mouse, ≈110 μg/kg for a 30 g mouse; injection performed by S. Pati), and experiments continued for ∼2 months. For histology, PRN cell counts were compared between control and DT-treated tissue. Diagonal paw-force coupling was assessed by calculating the Pearson correlation coefficient between the front-right (FR) and hind-left (HL) force traces within each trial. Trial-level correlation coefficients were compared between pooled controls and each ablation condition using independent-samples t-tests. Bonferroni correction was applied across the two comparisons (control versus DTR-Pan and control versus DTR-Glut), with statistical significance defined as an adjusted p < 0.05. These comparisons were exploratory because repeated trials were obtained from the same animals and each ablation group contained only two mice.

### Immunohistochemistry

Sections containing the PRN were stained with antibodies against tyrosine hydroxylase (TH) and serotonin (5-HT) to detect dopaminergic and serotonergic cells.

## Data and code availability

Code for hardware control and analysis is available at https://github.com/Egzona66/APA_setup.

## Acknowledgements

We thank the members of the Murray laboratory and the Sainsbury Wellcome Center for discussions and support.

## Author contributions

E.M. designed and built the behavioral paradigm, performed the behavioral, anatomical tracing and ablation experiments, and analyzed the data. F.C. acquired the Neuropixels recordings and developed the custom analysis code. A.J.M. supervised the work and acquired funding.

## Author declaration

All authors have read and approved the submitted version of the manuscript and consent to its posting on bioRxiv. This manuscript has not been accepted for publication and has not previously been published elsewhere.

